# Proteomic analysis reveals the recruitment of intrinsically disordered regions to stress granules

**DOI:** 10.1101/758599

**Authors:** Mang Zhu, Erich R. Kuechler, Joyce Zhang, Or Matalon, Benjamin Dubreuil, Analise Hofmann, Chris Loewen, Emmanuel D. Levy, Joerg Gsponer, Thibault Mayor

## Abstract

Heat-stress triggers the formation of condensates known as stress granules (SGs), which store non-translating mRNA and stalled translation initiation complexes. To gain a better understanding of SGs, we identified yeast proteins that sediment after heat-shock by mass spectrometry. Heat-regulated proteins are biased toward a subset of abundant proteins that are significantly enriched in intrinsically disordered regions (IDRs). SG localization of over 80 heat-regulated proteins was confirmed using microscopy, including 32 proteins that were not known previously to localize to SGs. We find that several IDRs are sufficient to mediate SG recruitment. Moreover, the diffusive exchange of IDRs within SGs, observed via FRAP, can be highly dynamic while other components remain immobile. Lastly, we showed that the IDR of the Ubp3 deubiquitinase is critical for SG formation. This work confirms that IDRs play an important role in cellular compartmentalization upon stress, can be sufficient for SG incorporation, can remain dynamic in vitrified SGs, and play a vital role during heat-stress.

**Summary:** The authors provide an in-depth proteomic study of yeast heat stress granule (SG) proteins. They identified intrinsic disordered regions (IDRs) as one of the main features shared by these proteins and demonstrated IDRs can be sufficient for SG recruitment.

## Introduction

Most organisms have a narrow temperature window for optimal growth and small thermal fluctuations may have detrimental effects (Richter et al., 2010). Maintaining protein homeostasis during temperature changes represents a major challenge to cellular systems as many polypeptides have only a marginally thermostable tertiary structure (Ghosh and Dill, 2010). Therefore, multiple cellular mechanisms are in place to respond to heat stress including, but not limited to, transcriptional up-regulation of heat shock proteins, inhibition of translation, and increased proteolysis. Part of the heat stress response in a wide range of cells from yeast and fly to mammals is also the formation of heat stress granules (SGs) (Farny et al., 2009; Groušl et al., 2009; Kedersha et al., 2005). SGs are membraneless ribonucleoprotein particles (RNPs) composed of mRNA and proteins. SGs assemble in synchrony with the stalling of protein translation (Buchan and Parker, 2009; Jain et al., 2016; Khong et al., 2017) and function as storage sites for non-translating RNA and stalled translation initiation complexes (Fan and Leung, 2016; Kimball et al., 2003). Studies have also shown other roles of SGs, such as the regulation of apoptosis (Arimoto et al., 2008; Eisinger-Mathason et al., 2008; Thedieck et al., 2013). Despite growing insights into composition and function of SGs, mechanisms of the formation, organization, and regulation of these membraneless compartments remains to be fully elucidated.

It is hypothesized that SG formation is governed by a demixing process known as liquid-liquid phase separation (LLPS) (Pak et al., 2016; Protter et al., 2018). Numerous recent studies have shown that several SG proteins form droplets or hydrogels *in vitro* (Hyman et al., 2014; Lin et al., 2015; Molliex et al., 2015; Patel et al., 2015; Riback et al., 2017). The rapid and diffusive exchange both within droplets and from protein-depleted to protein rich phases, which was measured *in vitro* by fluorescence recovery after photobleaching (FRAP), is consistent with the dynamics measured for SGs in several mammalian cells lines (Buchan and Parker, 2009). Intriguingly though, SGs in yeast appear to be different and display a non-amyloid solid-like state that is hexanediol resistant but SDS soluble (Kroschwald et al., 2015). Such a difference in material properties may be attributed to the presence of misfolded proteins in yeast SGs that reduce the mobility of other components (Cherkasov et al., 2013; Kroschwald et al., 2015). Supporting this notion, the deletion of the Hsp104 disaggregase delays SG dissolution during heat stress recovery (Kroschwald et al., 2015; 2018). It has also been proposed that SGs are constituted of a solid-like core surrounded by a more dynamic shell (Buchan and Parker, 2009; Jain et al., 2016), raising the question of whether individual components contained in yeast SGs remain mobile.

LLPS leading to the formation of RNP foci is often mediated by multivalent interactions between RNA and RNA-binding proteins as well as protein-protein interactions. These interactions can occur through folded domains, interaction motifs, and/or sequence patterns in intrinsically disordered regions (IDRs) that can be modified by post translational modifications (Elbaum-Garfinkle et al., 2015; Jonas and Izaurralde, 2013; Kedersha et al., 2016; Kroschwald et al., 2015; Nonhoff et al., 2007; Zhang et al., 2015). IDRs can enable promiscuous interactions in concert with specific domain interactions and, in some cases, have a more dominant effect; such as TIA-1 in SG formation and Lsm4 in yeast P-body formation (Decker et al., 2007; Gilks et al., 2004). Alternatively, other reports have shown that the presence of IDRs in proteins can weaken specific interactions (Protter et al., 2018). A comprehensive characterization of SG composition and the roles of IDRs in functional condensates will contribute to further understand SG regulation and their involvement in human diseases. Indeed, recent studies have shown that mutations of the SG protein FUS can affect its compartmentalization and lead to amyotrophic lateral sclerosis (ALS) or rare forms of frontotemporal lobar degeneration (FTLD) (Patel et al., 2015).

In this study, we identify a wide range of proteins that lose solubility upon heat stress and share common features, including many novel SG localizing proteins. We confirm that several of these proteins are not dynamic once recruited in heat SGs via FRAP. Furthermore, we demonstrate that IDRs alone can, in several cases, be sufficient for SG recruitment. We characterize the involvement of two IDRs to show that one can rapidly exchange in and out of the stress-induced foci, whereas the other is required for SG formation. This work sheds light on the importance of IDR in stress-induced protein compartmentalization.

## Results

### Global identification of proteins sedimenting after heat shock

Previously, we have shown that long proteins with low complexity IDRs are more pelletable in eukaryotic cells (yeast, HeLa, and mouse brain tissue cells) under unstressed conditions (Albu et al., 2015). In this study, we used a similar approach to delineate features shared among proteins that coalesce under stress conditions. We performed a systems-wide analysis of the *Saccharomyces cerevisiae* proteome by combining SILAC (Stable Isotope Labeling by Amino acids in Cell culture) with cell fractionation to identify proteins that sediment following an acute heat shock. Differentially labeled cells were either treated or not treated with heat shock at 45°C for 20 minutes, which induces a delay in growth without affecting viability (Fang et al., 2011). SILAC-labelled cells were subsequently mixed, lysed and fractionated by centrifugation before mass spectrometry analysis (Figure 1A). Among the 3734 proteins identified in the supernatant fraction with at least 2 peptides, 2717 proteins were quantified in three independent experiments (Figure 1B; Table S1). The analysis showed that more than 60% of the quantified proteins remained predominately in the supernatant post heat shock. In contrast, a subset of 165 proteins showed a 90% or higher depletion from the supernatant after heat shock. Ola1 had the highest quantified depletion; with less than 2% of Ola1 remaining in the supernatant after heat shock. In parallel, we identified 3528 proteins from the pellet fraction, among which 2558 proteins were quantified in all three experiments (Figure 1C; Table S2). Most proteins that were quantified in both the supernatant and pellet fractions displayed inversely correlated ratios consistent with the theoretical curve (Figure 1D). However, a small group of proteins were depleted from both fractions after heat shock, possibly indicating these proteins are depleted from the cell after the stress. Protein localization analysis showed that this small group of proteins mostly localize in the nucleus or nucleolus (Table S3). Analysis of the total cell lysate showed that levels of these proteins remained mostly unchanged after heat shock (Figure S1A). Therefore, these proteins most likely form compact structures or macro-molecular assemblies that were depleted following the removal of cell debris performed at low speed centrifugation. In order to circumvent this issue and not omit data from highly pelletable structures, we focused our analysis on proteins depleted from the supernatant.

**Figure 1.**
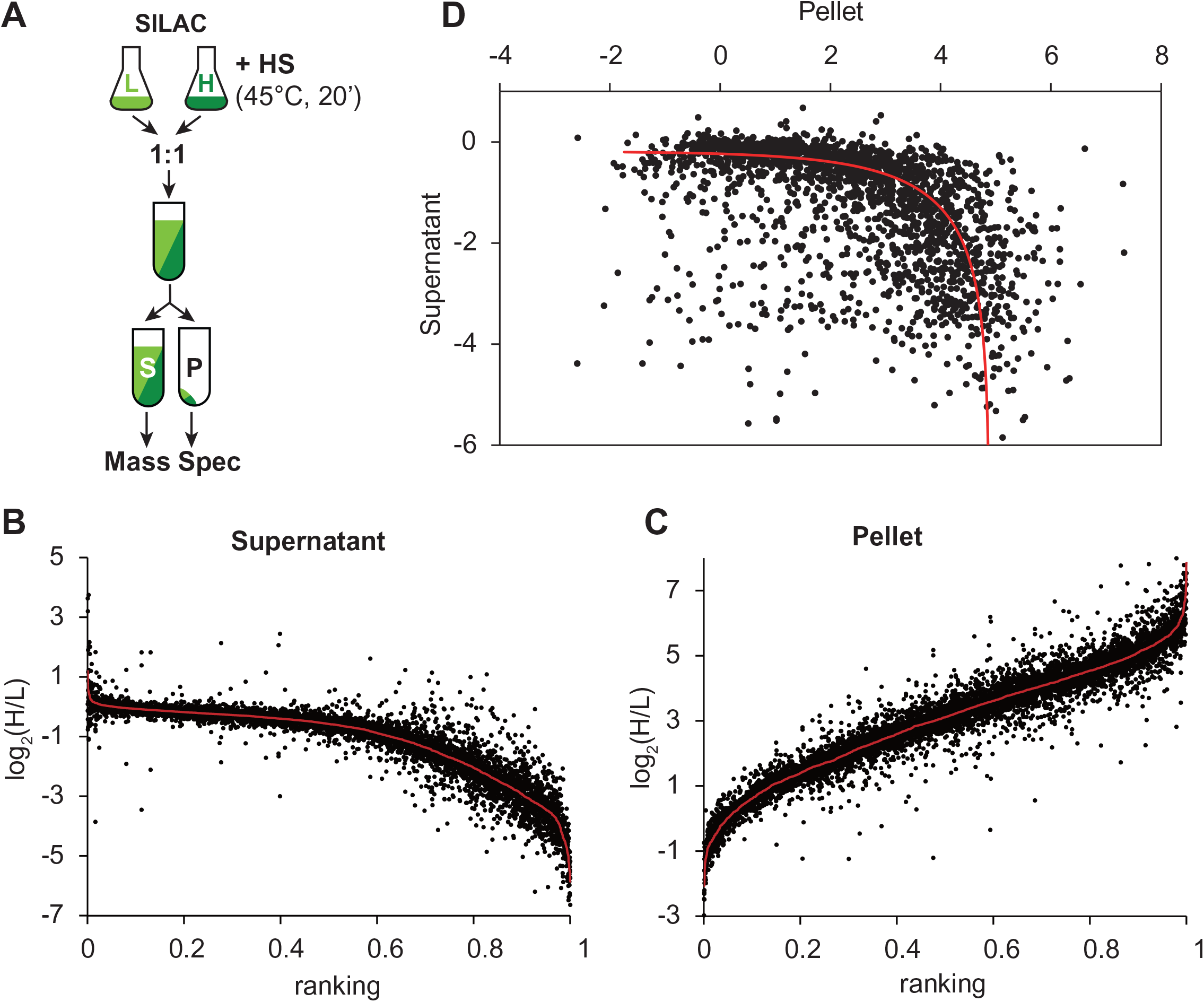
Identification of pelletable proteins after heat shock by mass spectrometry. (A) Schematic of the SILAC mass spectrometry approach to identify pelletable proteins after heat shock (HS). (B and C) log_2_ratios (H/L) of proteins after and before heat shock in supernatant (B) and pellet (C) fractions. Proteins are ranked based on descending ratios, each black dot represents one of three analyses, and the red line marks the geometric mean of three replicates. (D) Plot of the averaged log_2_ (H/L) of 2060 proteins quantified in both the pellet and supernatant fractions. Red line indicated the theoretical curve.

Next, we sought to evaluate how the data resulting from our supernatant analysis compared to similar recent studies. Many of the proteins that have a ten-fold or more depletion from the supernatant (165) overlap with proteins known to aggregate or form cellular inclusions in previous studies (Figure S1B) (Cherkasov et al., 2015; Wallace et al., 2015). Some of these proteins (9) were classified as super-aggregators by Drummond and colleagues (Wallace et al., 2015) and many of these proteins (38) were shown to co-localize with the Pab1 stress granule marker after heat shock by Bukau and colleagues (Cherkasov et al., 2015). Gene ontology (GO) analysis also showed that this group of proteins highly depleted from the supernatant is enriched for cytosolic SG proteins (p-value: 3.19e-13; count: 22). To further evaluate our results, we also sampled 13 candidates among the cytosolic proteins quantified by mass spectrometry to assess their localization in the cell when tagged to GFP. Proteins that were not found to be affected by heat shock during the proteomic analysis remained diffuse within the cell, while proteins that became depleted from the supernatant fraction displayed distinct foci (Figure S1C), indicating that many proteins strongly depleted from the supernatant are likely recruited to foci after acute heat shock. Importantly, most chaperone or co-chaperone proteins were found to remain in the supernatant fraction (Figure S1D), presumably while associated to misfolded proteins that do not aggregate. Surprisingly, Sis1 Hsp40 remained largely in the supernatant, despite having been shown to be recruited to amyloidic-site deposits (Park et al., 2013). In contrast, small heat shock proteins Hsp26 and Hsp42 were depleted, consistent with their reported association to stress-induced foci (Escusa-Toret et al., 2013; Specht et al., 2011). Additionally, many translation initiation factors and RACs (ribosome associated chaperones) were depleted from the supernatant (Figure S1D), consistent with previous studies (Cherkasov et al., 2015; Wallace et al., 2015). Groušl *et al*. showed that proteins from 40S ribosomal subunits were recruited into SG upon heat shock (Groušl et al., 2009). However, it was later shown that ribosomal proteins do not localize in yeast SGs (Cherkasov et al., 2013). In agreement with the latter report, all cytosolic ribosomal proteins identified remained largely in the supernatant after heat shock with the exception of Rps31. In contrast, a majority of mitochondrial ribosomal proteins were found to be depleted by over 50% from the supernatant, indicating that these complexes may aggregate within the mitochondria.

### Heat inducted pelletable proteins share common characteristics

To identify potential causes for the increased “pelletability” (i.e. the ability to be enriched in the pellet after centrifugation) upon heat shock, we mapped features that are shared among proteins that mostly remained in the supernatant (hereafter called soluble; 1,704 proteins) versus proteins that were mostly depleted from the supernatant and referred as pelletable (1,013 proteins; Figure 2A). The pelletable proteins were significantly longer (Figure 2B; n and p-values are reported in Table S4) and were overall less hydrophobic (Figure 2C). In agreement, these pelletable proteins also comprised a greater proportion of charged, both positively and negatively, residues (Figure S2A-C). Additionally, these proteins were predicted to be more disordered (Figure 2D). In good agreement with these last three observations, pelletable proteins contained more solvent exposed residues (Figure 2E). These results suggest that these proteins have more expanded structures with flexible IDRs that could facilitate transient interactions with other proteins or nucleic acids that may facilitate their enrichment in the pellet fraction.

**Figure 2.**
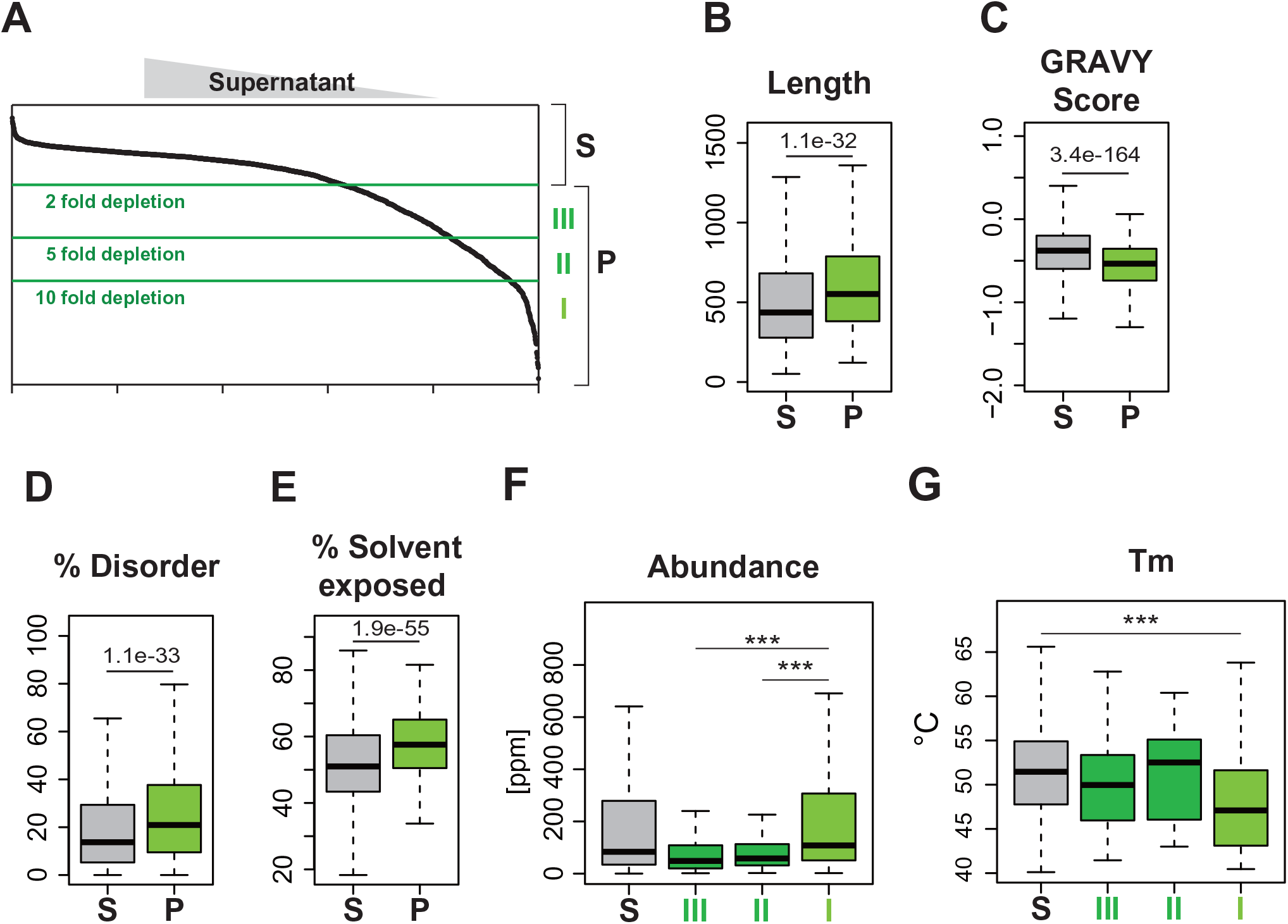
Protein feature analysis of pelletable proteins upon heat shock. (A) Binning of data using 2, 5, 10-fold depletion from the supernatant as cutoff. (B-E) Box plots comparing distributions of soluble (S) and pelletable (P) proteins. The log_2_(H/L) in the proteomic analysis of the supernatant fraction was > −1 and < −1 for soluble and pelletable proteins, respectively. The following analyses are shown for sequence length in number of amino acids (B), hydrophobicity based on GRAVY score (C), % disordered (D), % solvent exposed (E). p-values are shown. (F-G) Box plots comparing distributions of proteins in designated bins for protein abundance expressed in part per million (ppm) (F) and T_m_ (°C) (G). All n and p-values are reported in Table S4.

As proteins that were highly depleted from supernatant after heat shock formed more distinct foci (Figure S1C) and were enriched for SG proteins, we decided to further separate the pelletable proteins into three additional groups based on their fold depletion: group I) 165 proteins depleted over 10-fold; group II) 287 proteins depleted between 5 and 10-fold, and group III) 561 proteins depleted between 2 and 5-fold; Figure 2A). Upon initial examination of these groups, we found that group I was uniquely enriched for RGG motifs (25 proteins, p-value: 7.2e-6), which are known to aid in the formation of biological LLPS media and are implicated in SG formation (Chong et al., 2018). Surprisingly, proteins displaying over a 10-fold depletion from the supernatant (group I) were more abundant in comparison to other pelletable proteins (groups II and III; Figure 2F). In addition, only proteins in that group displayed a significantly lower apparent melting temperatures (T_m_) in comparison to soluble proteins (Figure 2G). In contrast, all the previously described features associated to pelletable proteins were still enriched in the three subgroups in comparison to soluble proteins (Figure S2D-H). Given that proteins in group I display higher protein abundances coupled with lower apparent melting temperatures suggests that protein pelletability is likely driven by an intrinsic function to regulate protein coalescence during cellular stress response induced by heat shock.

### Heat induced pelletable proteins are enriched for stress granule localized proteins

Given the high GO enrichment for SG localization among the proteins that were mostly depleted from the supernatant after heat shock, we used confocal microscopy to determine which proteins co-localized with the poly-A binding protein and SG marker Pab1. Specifically, we assessed the 136 strains that were available from the Yeast GFP collection for which the proteins were found depleted over 10-fold from the supernatant (excluding membrane proteins). Following mating with a strain containing Pab1-mCherry, microscopy analysis identified 86 GFP tagged proteins with Pearson correlation coefficient (PCC) for co-localization with Pab1 larger than 0.2 after heat shock (Figure 3A; Table S5). Notably, a drop in the PCC followed this cut off, and manual validation of microscopy images confirmed that no co-localization with Pab1 could be observed in these cells (Table S5). Typically, proteins with high (>0.45) and mid-range (<0.45 and > 0.2) PCC displayed either full or partial colocalization with Pab1, respectively (Figure 3B, C; Table S5). To assess the effect of the fluorescence tag on possible partial co-localization with Pab1, the GFP and mCherry tags were switched on a subset of these proposed heat SG components. In contrast to previous analysis, three tested proteins tagged to mCherry showed a full colocalization with Pab1-GFP after heat shock (Figure S3A). Differences in the results (partial versus full co-localization of the assessed proteins with Pab1) can be attributed to the different tagging or marginal variations in the methods used to assess co-localization. Notably, a longer delay between the heat shock and cell imaging was introduced when assessing Pab1-mCherry co-localization due to technical limitations. Consistent with our previous analysis, heat-induced SG proteins have a lower apparent T_m_(Figure 3D). Interestingly, SG proteins with higher PCC for Pab1 co-localization displayed slightly lower apparent T_m_ (difference not significant). Moreover, specifically SG proteins with higher PCC were more often found to be associated to RNA (Figure 3E). Of note, the lower T_m_ is not characteristic of RNA binding proteins, as RNA-binding proteins did not display lower apparent T_m_ values (Figure 3D). Moreover, SG proteins with higher PCC values had a greater number of identified phosphorylation sites among known phosphorylated proteins (Figure S3B). These results indicate that phosphorylation may play an important role in regulating SGs, as previously indicated (Reineke et al., 2012; Kedersha et al., 1999; Yoon et al., 2010; Reineke et al., 2017). Low complexity regions (LCR) have been implicated in LLPS (Mitrea and Kriwacki, 2016; Molliex et al., 2015). We found that among proteins that co-localize with Pab1 there was a significant enrichment for LCR with specific amino acids, especially for SG proteins with high PCC (Figure S3C). To summarize, we confirmed that large portion of proteins depleted from supernatant are recruited into heat-induced Pab1-containing granules and identified 32 novel SG proteins that were not identified as such in previous studies (Cherkasov et al., 2015; Wallace et al., 2015).

**Figure 3.**
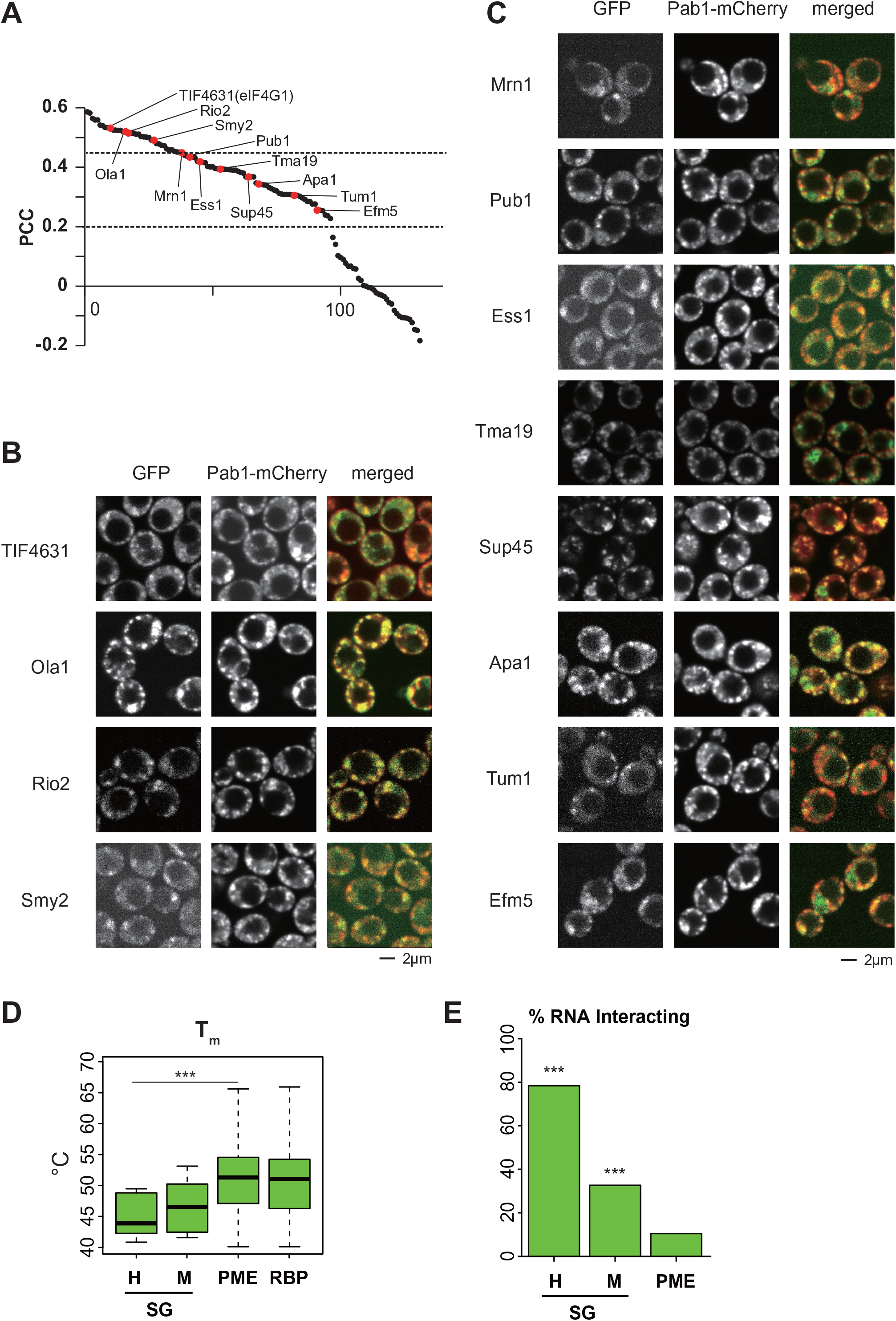
Co-localization analysis with stress granule marker Pab1. (A) Pearson correlation coefficient (PCC) of the colocalization with Pab1-mCherry with 136 GFP-tagged proteins assessed by confocal microscopy in diploid cells and ranked in descending order. (B-C) Representative images of quantified cells of a few selected GFP-tagged proteins (designated in A) with high PCC (B) or median PCC (C) values. (D) Box plot of the distributions of the apparent T_m_ among SG proteins with high PCC values (H; n= 15) and mid-range PCC values (M; n = 9), the proteome (PME; n=706), and all RNA-binding proteins in the proteome (RBP, n = 244). (E) Bar plot shows whether SG proteins with high or mid-range PCC were found to bind RNA or not. All n and p-values are reported in Table S4.

### Proteins in yeast heat stress granules are largely immobile

It was previously shown that yeast SGs display a solid-like behavior, contrasting the liquid-like SG found in mammalian cells (Kroschwald et al., 2015). To validate these findings, we directly measured the mobility of several proteins sequestered to SGs in yeast cells using fluorescent recovery after photobleaching (FRAP). We selected the SG marker Pab1, the Ola1 ATPase implicated in read through of premature stop codons (high PCC in Figure 3A), and the diadenosine phosphorylase Apa1 (mid-range PCC). After 10 minutes of heat shock at 45°C, the Pab1 signal in the photobleached foci recovered to about 30% of the initial signal intensity while the fluorescent signal of not photobleached foci decreased by about 20% (Figure 4A). The recovery of Pab1 was reduced to 21-22% after a 15 and 20 minutes heat shock. Noticeably, despite the recovery of some Pab1 signal, no distinct inclusion site was recovered in the photobleached area. Interestingly, when FRAP was performed before the formation of distinguishable Pab1 foci (after a 5-minute heat shock at 45°C), the cytosolic region revealed decreased mobility (Figure S4A). We used Tpi1, a protein which remains soluble after heat shock based on our mass spectrometry data, as a control. In agreement, Tpi1-GFP did not form foci after 15 minutes of heat shock at 45°C (Figure S4B). Importantly, the signal in the photobleached area in heat shocked Tpi1-GFP cells showed a fast recovery that was below the 5 second time frame required for image acquisition (Figure S4B). These results indicate that, whereas a smaller fraction of Pab1 may stay dynamic or start to dissociate, most of Pab1 remains associated to the foci, similar to what was observed in Pab1 droplets assembled *in vitro* (Riback et al., 2017). Additionally, Ola1 showed much lower recovery (Figure 4B). Ola1 formed detectable foci after 5 minutes of heat shock at 45°C. After photobleaching, a minute increase of signal intensity was observed with a recovery of approximately 15% after 90 seconds. Longer periods of stress prior to photobleaching (10 and 15min heat shock) led to a further decreased recovery of 4-7% of the initial signal intensity. Remarkably, Apa1 was not more dynamic in SG despite displaying a lower PCC with Pab1 (Figure 4C). Apa1 showed a 9-10% recovery after 10 and 15 minutes heat shock at 45°C. These results confirm that components of yeast heat SGs display poor protein dynamics once recruited to these heat-induced compartments.

**Figure 4.**
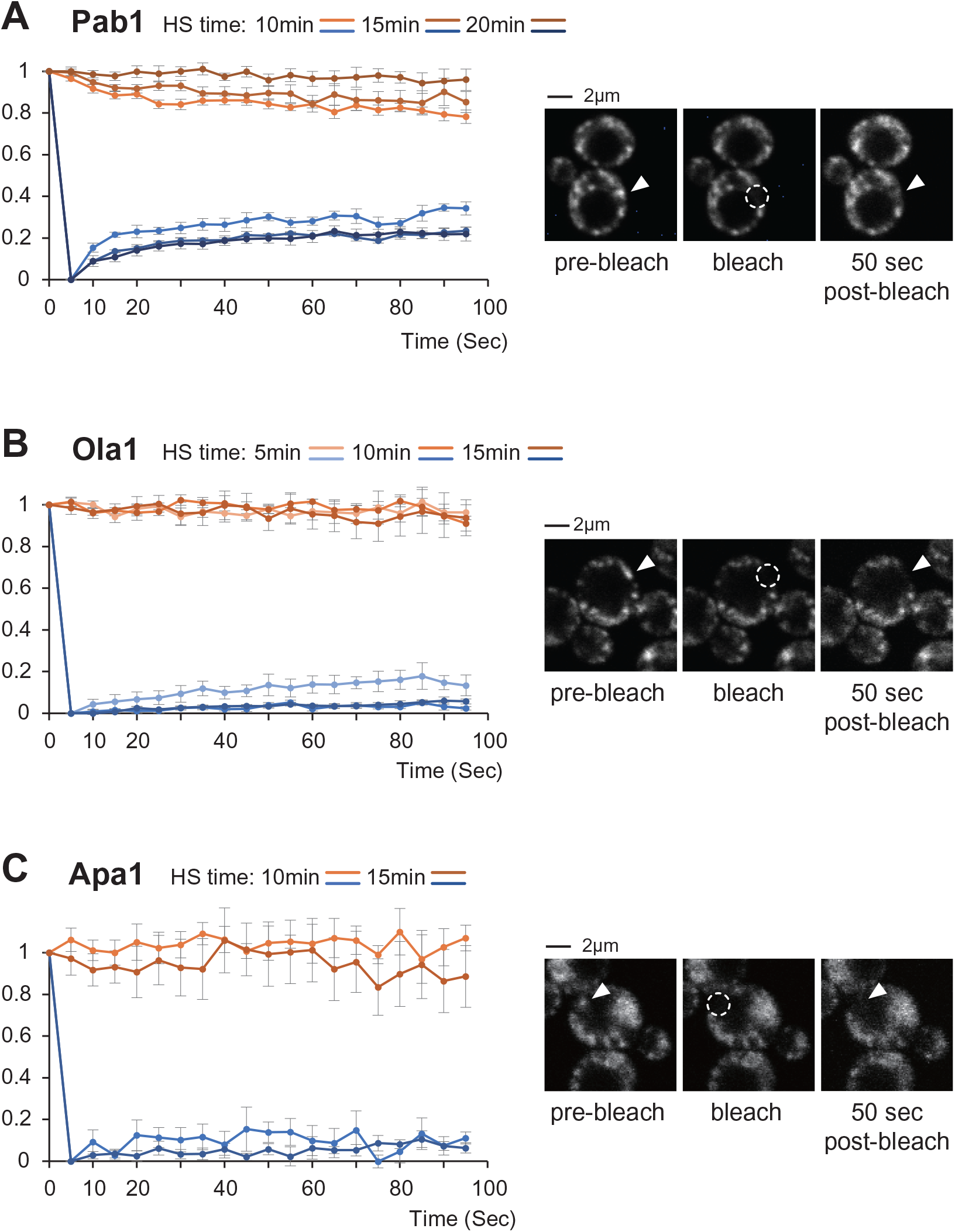
Components of heat-induced stress granules are poorly mobile. **(A-C)** Fluorescent Recovery After Photobleaching (FRAP) and Fluorescence loss in photobleaching (FLIP) of Pab1 (A), Ola1 (B) and Apa1 (C) tagged to GFP at their endogenous locus. The graphs show the averaged signal intensities from 6 replicates (with Std errors) in the indicated time points after incubating the cells at 45°C for the indicated time periods. FLIP data was measured in three ROI containing unbleached foci within a bleached cell. Representative cells are shown on the right. The dotted circles represent the photobleached ROIs and the arrowheads indicate the analyzed foci before and 50 seconds after photobleaching.

### Intrinsiclly disordered regions can be sufficient for stress granule recruitment

Our finding that proteins with reduced solubility after heat stress are enriched in IDRs and localize to SGs is consistent with the observation that ribonucleoprotein particle (RNP) proteins often contain IDRs (Decker et al., 2007; Hennig et al., 2015; Jain et al., 2016; Protter et al., 2018). Next, we sought to evaluate whether IDRs are necessary or sufficient for proteins to coalescence into heat SGs. It has previously been shown that a short, disordered segment of low sequence complexity is responsible for the recruitment of Cdc19 to starvation-induced granules, and that this short peptide is sufficient to drive GFP into granules (Saad et al., 2017). We employed a similar approach and expressed IDRs of different SG proteins fused to GFP from a plasmid. Five of the fourteen IDRs that we assessed were able to drive GFP localization into Pab1 foci after heat shock while the signal remained diffused in untreated cells (Figure 5A; Figure S5A, B). IDRs of Ubp3, Psp2, Dcp2, and Tif4631/eIF4G1 showed strong recruitment to foci, whereas the IDR of Mot2 led to a weaker but distinct GFP signal in Pab1 foci. IDRs of Ubp3, Dcp2, Mot2, and Tif4631 contain one or more patches of predicted aggregation-prone regions or with enhanced score for prion-like amino acid composition (Figure 5A). In addition, all five IDRs contain regions with low complexity amino acid sequences. However, those low complexity regions lack a consensus in amino acid enrichment. Ubp3, Psp2, Tif4631 and Dcp2 IDRs share low complexity regions (LCRs) enriched with in asparagine only excluding the Mot2 SG incorporated IDR without this feature, however asparagine LCRs are found in three of the nonSG recruited IDRs as well (Smy2, Tif4631, and Caf40; Figure S5C). Similarly, proline IDRs were favored in SG included IDRs, appearing on six separate occasions across four different proteins while only appearing four times across the nine non-coalescing IDR segments. Histidine did uniquely appear in those IDRs which are recruited into SGs, however this region only appeared in Ubp3. Additionally, all five IDRs contain molecular recognition features (MoRFs), which are short segments in IDRs that adopt secondary structures upon binding to other domains. Nonetheless, IDRs that failed to be recruited to SGs also contain similar features (Figure S5C). Thus, none of these features may be sufficient, when examined in the general case, to explain IDR SG incorporation. Next, we sought to determine whether these five IDRs could also be recruited to SGs induced by other stresses. Excluding the IDR of Tif4631, these IDRs were also recruited into Pab1-containing granules under glucose starvation (Figure 5B, Figure S5D). Similarly, the IDR of Ubp3 and Psp2 were distinctively recruited to Pab1 foci upon azide treatment (Figure 5C), while the recruitment of the IDR of Mot2 and Dcp2 appeared more modest. Importantly, these results show that several IDRs could directly be recruited into SG, in contrast to a recent report in which none of the assessed IDR was sufficient to target components to RNP granules (Protter et al., 2018).

**Figure 5.**
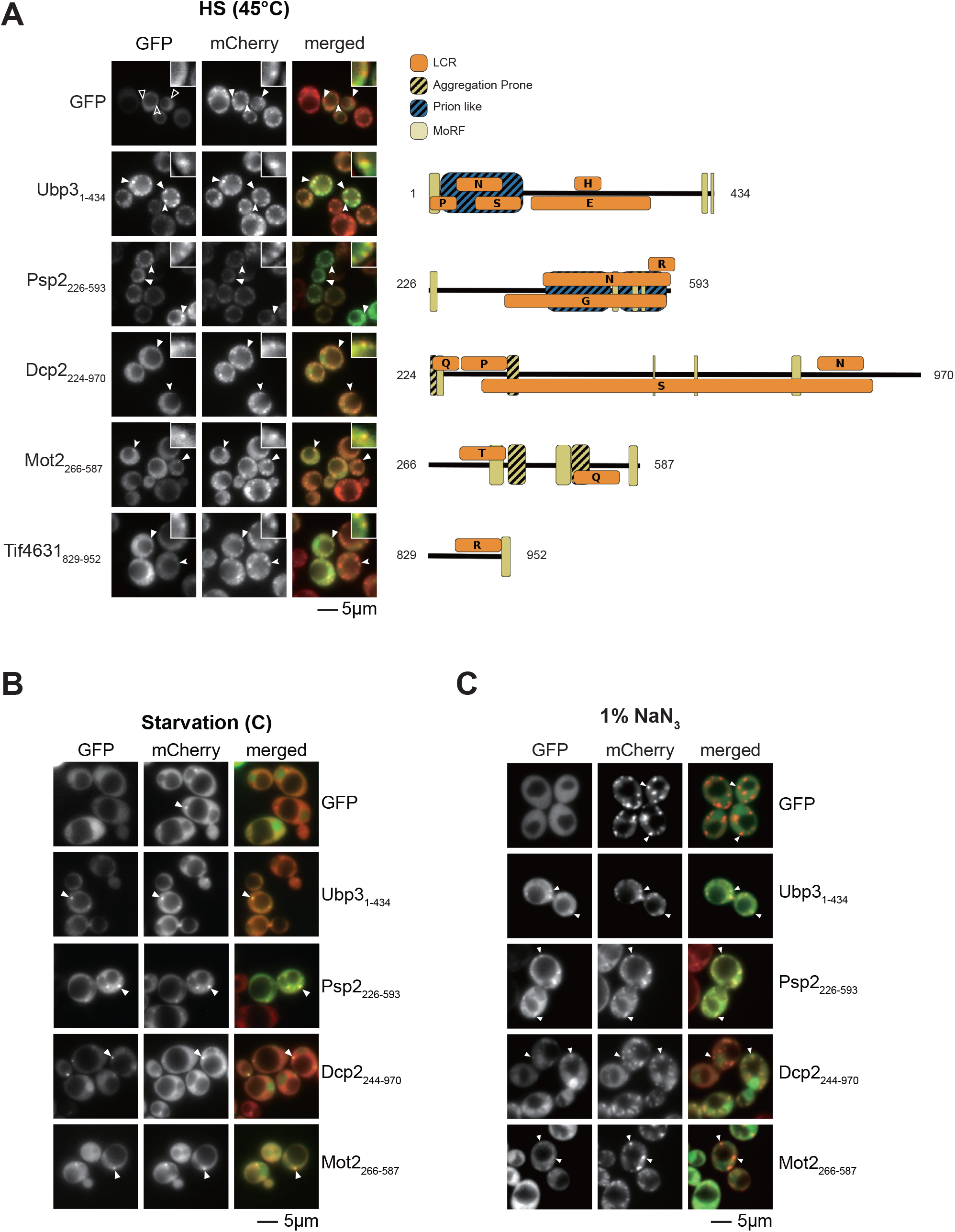
Recruitment into stress granules by intrinsic disorder region. (A) Representative images of Pab1-mCherry cells that expressed from a plasmid the designated disordered regions tagged with GFP. Cells were exposed to a 15 min heat shock at 45°C prior to imaging. Filled arrowheads designate representative Pab1 and GFP foci. The insets show a five-times magnified image of the foci designated by a sunken arrowhead. Feature analysis of the corresponding IDR is shown on the right side. Orange boxes indicate low complexity regions with letters to indicate specific amino acid enrichment, black striped boxes indicate predicted aggregation prone regions, blue striped boxes indicate prion-like composition, and yellow boxes indicate predicted MoRFs. Only highly significant (p ≤ 1e-6) low complexity regions are pictured. (B-C) Same as (A) but upon carbon starvation for 90min (B) or after 30min incubation at 30°C with 1% NaN_3_ (C). Exposure time for GFP was increased in (C) due to dimmer signal in these conditions.

### Association of IDRs to SG compartments display different properties

Ubp3 is a deubiquitinase and was recently shown to be important for the assembly of SGs upon starvation and heat shock (Nostramo et al., 2016). Ubp3 was also shown to be involved in Rsp5-dependent proteasome degradation of misfolded proteins upon heat shock (Fang et al., 2016). Psp2 is asparagine-rich RNA-binding protein that was recently shown to promote the formation of P-bodies in some conditions (Rao and Parker, 2017). The IDR of Psp2 also contains several RGG motifs in its IDR, which have previously been implicated in protein LLPS (Chong et al., 2018). Interestingly, while the IDR of Ubp3 remained in the pellet fraction after centrifugation, the IDR of Psp2 was largely in the supernatant after heat shock (Figure 6A). These findings suggest that the association of Psp2’s IDR to the SG is potentially more transient allowing for rapid exchange between the pelletable and supernatant fractions. To test this hypothesis, FRAP was employed to compare the mobility of both IDRs. As expected, we found that Ubp3’s IDR displayed minor recovery; similar to Ola1, Apa1, and full length Ubp3 (Figure 6B). In contrast, the IDR of Psp2 was found to be rapidly recruited to the foci within 5 seconds after photobleaching with a signal recovery of ∼80% of the initial intensity (Figure 6B). This dynamic recovery was limited to Psp2’s IDR, as the full length Psp2 only showed modest resurgence. These results suggest that the IDRs of Ubp3 and Psp2 associate to SGs using different mechanisms. To determine whether these interactions are intra-or inter-molecular, we next assessed the ability of these IDRs to be recruited into foci in absence of the respective full-length proteins. The IDR of Ubp3 but not of Psp2 coalesced in *ubp3Δ* and *psp2Δ* cells, respectively (Figure 6C), suggesting that the IDR of Ubp3 may be recruited to the heat SG upon binding to another protein. In contrast, as the IDR of Psp2 requires the presence of the full length Psp2 for SG localization, the IDR likely interacts with the N-terminal region of Psp2. Indeed, the structured region of Psp2 (residues 1-225), which has one RNA recognition motif (RRM), was recruited into Pab1 foci independently of the IDR (Figure S6A). These results show that IDRs associate to SG in different manners, as the IDR of Psp2 appears more dynamic. Particularly, these results indicate that the yeast heat-induced compartments are not static macro-assemblies with limited diffusion, as some of its components can freely associate and dissociate following their formation.

**Figure 6.**
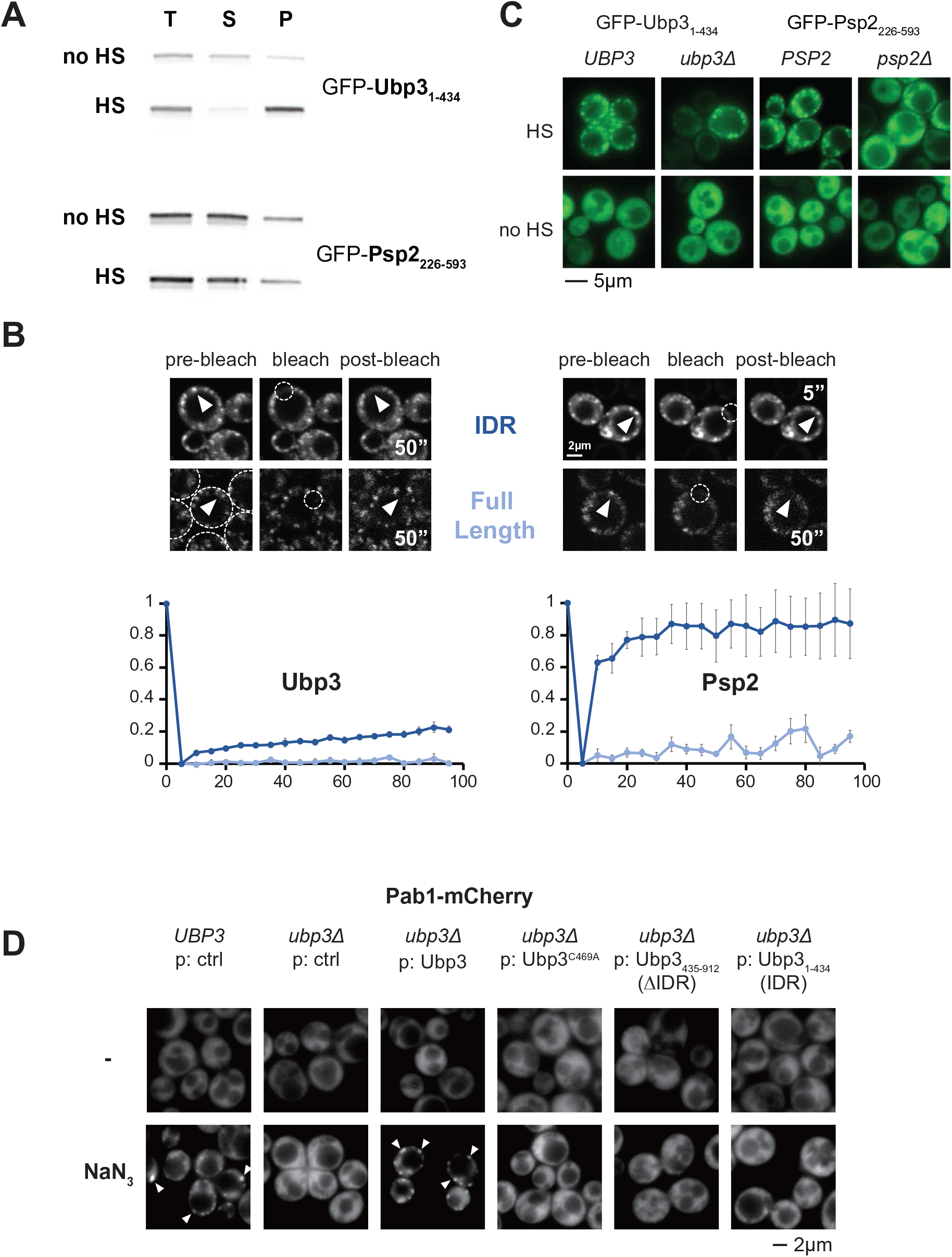
IDRs from Ubp3 and Psp2 display different properties. (A) Western blot against GFP in the indicated lysate fractions derived from unstressed (no HS) and heat shocked (HS) cells. T: Total cell lysate, S: supernatant, P: Pellet. (B) FRAP in heat shocked cells (15 min at 45°C) expressing the designated disordered regions (dark blue) or full-length proteins (light blue) tagged with GFP. The IDR segments fused to GFP were expressed from a plasmid, whereas the full-length proteins were endogenously tagged. Representative cells are shown on the top. The dotted circles represent the photobleached ROIs and the arrowheads indicate the analyzed foci before and 50” or 5’’ after photobleaching. The graphs show the averaged signal intensities from 6 cells quantified sequentially (with Std errors) in the indicated time points. (C) Representative images of the indicated wildtype or mutant cells expressing from a plasmid the designated disordered regions tagged with GFP before and after heat shock. (D) Representative images of Pab1-mCherry cells with the indicated genetic background and protein expressed from a plasmid (p) before (-) and after 30min incubation at 30°C with 1% NaN_3_. Arrowheads designate representative Pab1 foci. No Pab1 inclusion was observed in the absence of wild-type Ubp3 in three independent experiments. Of note, the fragment of Ubp3 that lacks the IDR also formed foci in azide treated cells (data not shown).

### The IDR of Ubp3 is required for its role in stress response

We next wanted to assess whether the IDR of Ubp3 was also important in SG formation. While we were not able to observe a clear decrease of SG formation in *ubp3Δ* cells by monitoring Pab1-foci under heat shock condition (Figure S6B), lack of Ubp3 led to the absence of Pab1 foci in azide treated cells (Figure 6D), as previously reported (Nostramo et al., 2016). Importantly, expression of the truncated Ubp3 that lacked the IDR could not rescue SG formation (Figure 6D). Intriguingly, the loss of Ubp3 function also led to a marked increase of cells that do not display a singular and large vacuole after heat shock (Figure S6B). Accordingly, the expression of wild-type Ubp3 abrogated this phenotype in *ubp3Δ* cells, whereas the expression of the IDR alone led to a partial rescue and the expression of the truncated Ubp3 that lacked the IDR only a mild rescue (Figure S6B). These results highlight how the presence of Ubp3 IDR plays a role in the stress response and the assembly of SG in some conditions, possibly by recruiting substrates and the Bre5 cofactor, which forms a complex with Ubp3 by interacting with the Bre5 binding domain within the IDR of Ubp3 and is required for the activity of the deubiquitinase (Li et al., 2005; 2007).

## Discussion

In this study, we employed a proteomic approach to assess how heat shock alters protein sedimentation and show that heat-induced pelletable proteins share common characteristics. Combining proteomic results with high-content imaging, we identify 32 novel heat SG-localizing proteins. We further confirm that several components of yeast heat SGs are highly static and that IDRs are able to drive SG recruitment.

While a main focus of this study is the recruitment of proteins into cytosolic heat SGs, several proteins depleted from the supernatant fraction after heat shock do not localize to Pab1-containing inclusions. These include proteins from mitochondria, the nucleus and nucleolus, similar to what was observed previously (Cherkasov et al., 2015). Interestingly, almost all of the proteins that were lost in heat shocked cells in the low centrifugation pre-clearing step prior to fractionation localize to the nucleus or nucleolus, and more than one-third of those proteins are involved in the pre-90s ribosome assembly. As nucleolus proteins phase separate during ribosome assembly (Mitrea et al., 2018), it is possible that these macro-assemblies in the nucleolus undergo additional rearrangements upon heat shock leading to a higher density compartmentalization. We found that mitochondrial proteins also form foci after heat shock (Table S5). Thus, mitochondria or some elements of this organelle may also undergo a drastic rearrangement during acute heat stress.

Previous work from Drummond and colleagues indicates that protein coalescence after heat shock is a reversible process that does not rely on proteolysis to “regenerate” the bulk of soluble proteins after the stress recovery (Wallace et al., 2015). We found that proteins that were the most depleted from the supernatant (group I) were also highly abundant. These observations are consistent with the work from Drummond and colleagues, as the degradation of abundant proteins upon stress recovery would present a clear evolutionary disadvantage. The fact that these abundant proteins are mostly depleted from the supernatant fraction (>90%) also indicates that the compartmentalization of these proteins is highly efficient. Recent proteomic studies using proximity tagging showed that many stress granules components interact with each other in both unstressed and stressed cells (Markmiller et al., 2018; Youn et al., 2018). These studies indicate that the process is likely tightly regulated in order to only drive coalescence after the stress. The prevalence of charged residues and potential phosphorylation sites among pelletable and SG proteins support a role for electrostatic interactions in stress-induced condensates that may be, in part, regulated by kinases.

SGs that assemble upon an acute heat shock in yeast have been shown to harbor misfolded proteins (Cherkasov et al., 2013; Mateju et al., 2017). The lower apparent melting temperature of SG proteins suggests that many SG proteins undergo a possible structural transition under heat shock. Such a change may allow for the exposure of buried residues for additional promiscuous, hydrophobic interactions that would increase interaction valency thus aiding the demixing process of heat SG components (Li et al., 2012). It should be noted that the apparent T_m_ measurements derived from the proteomic analysis were done at the peptide level, and that the measured values were not always equivalent across a given polypeptide (especially in multi domain proteins) (Leuenberger et al., 2017). Therefore, misfolding of highly pelletable proteins could be partial (i.e. specific to a domain within a polypeptide) to potentially retain functionality of the protein as shown before (Wallace et al., 2015) and to facilitate recovery. In addition to proteins that are predominantly depleted from the supernatant after heat shock, a sizeable portion of the proteome was also affected (designated groups II and III in Figure 2). Examination of a few of these proteins showed that they were weakly recruited to foci (e.g., YLR287c and Dph6 in Figure S1C). One possibility is that many of these proteins misfold, leading then to SG recruitment. Interestingly, these proteins are of lower abundance in the cell, consistent with a possible lower selective pressure to maintain them folded upon acute stress. It would be interesting to determine whether newly synthesized proteins may also be more susceptible to misfolding and SG recruitment.

We found that there is an enrichment of IDRs among the heat-induced pelletable proteins. Many studies have investigated the role of IDRs particularly prion-like domains and IDRs with low complexity in protein phase separation (Decker et al., 2007; Gilks et al., 2004; Kato et al., 2012; Protter et al., 2018). In contrast to some of these previous studies, we found at least five IDRs that are sufficient to mediate SG recruitment of the fused GFP moiety. All five IDRs contain distinct protein sequence features that may contribute to their ability for SG recruitment but none that are universal or unique. Perhaps the presence of longer prion-like domains in the IDRs of Psp2 and Ubp3 contribute to a higher likelihood to LLPS, and thus to SG incorporation. All of the tested IDRs contain several compositionally biased regions of low complexity, however due to the relatively small group of IDRs assessed, it remains difficult conclusively link any of these features to SG recruitment with confidence. Moreover, we also speculate that it is very likely that SG incorporation is either controlled through direct interactions or through the fine tuning of protein solvation behavior which, by necessity, is specific to each protein. Therefore, IDRs recruited to SGs may not share common features. Nonetheless, the analysis of the diffuse behavior of Ubp3 and Psp2 in SGs revealed that IDRs located to SGs can display pronounced differences in dynamics, which is presumably reflective of their respective affinity for other domains within these membraneless compartments. As the structured regions of both Ubp3 and Psp2 proteins were also recruited to Pab1 foci (Figure S6A), their IDRs may further facilitate recruitment to SGs by adding valency and/or increase the affinity for partners within the stress compartment. In the case of Ubp3, the IDR also plays a role in SG assembly. Whereas presence of IDR is a prominent feature among SG proteins, many IDRs were not directly recruited to SG. In some cases, their affinity to other SG elements alone may be too weak. Alternatively, they could also potentially facilitate recovery after the stress, similar to what recently shown for the Sup35 prion-like domain (Franzmann et al., 2018).

## Methods

### Yeast strains and plasmids

For SILAC, a modified BY4741 *Saccharomyces cerevisiae* yeast strain was used (YTM1173, *MATa*, *his3Δ1*, *leu2Δ0*, *ura3Δ0*, *MET15*, *arg4Δ::KanMX6*, *lys2Δ0*). The Pab1-mCherry strain (YTM1920; *MATα, his3Δ1, leu2Δ0, lys2Δ0, ura3Δ0, Pab1-mCherry::KanMX6*) was generated by homologous recombination using a yeast codon-optimized mCherry placed into pFA6a-kanMX6 (BPM866). *TMA1*, *ESS1* and *MRN1* were similarly modified from the BY4742 strain. Cells with mCherry tagged alleles were then mated with selected strains from yeast GFP collection (Huh et al., 2003) to generate diploid cells for colocalization analysis. For the analysis of IDRs, as well as Ubp3 and its mutants, the indicated gene segments were amplified from yeast genomic DNA and subloned C-terminally of GFP using EcoRI/NotI and SalI restrictions sites, into a modified pRS313 with the GPD promoter and *PGK1* terminator sequences (BPM1174-1220, 1453-1454).

### Yeast culture and Sample preparation

Yeast cells were incubated at 25°C, unless otherwise stated. For SILAC, cells were grown in light labeled (0.03mg/ml Lys0, 0.02mg/ml Arg0) or heavy labeled (0.03mg/ml Lys4, 0.02mg/ml Arg6) amino acid in synthetic defined medium with 2% dextrose for at least 7 generations. Heat shock was performed in a shaking water bath at 45°C for 20 minutes before harvest. Equal OD_600_ of cells from both labels were collected at mid log phase (OD_600_=0.8-1) by centrifugation at 3,220x*g* for 5 minutes at 4°C, washed twice with cold 1×TBS (50mM Tris pH 7.5, 150mM NaCl), mixed, and re-suspended in equal volume of 2×Native lysis buffer (200mM Tris pH 7.5, 150mM NaCl, 2mM PMSF, 2×Protease Inhibitor Cocktails (Sigma-Aldrich), 2mM 1, 10-Phenanthroline). Re-suspended cells were snap-frozen drop by drop in liquid nitrogen. The resulting cells pellets were lysed by cyro-grinding in liquid nitrogen with a mortar and pestle mounted on an electric driver (OPS Diagnostics). The lysate was thawed on ice and further diluted 3 times by adding ice-cold 1×Native lysis buffer and 1% NP40 (final concentration). Total cell lysate was pre-cleared twice by centrifugation at 1,000×*g* at 4°C for 15min. Pellet fraction was separated by centrifugation at 16,100×*g* at 4°C for 15min. The insoluble protein pellet was washed twice with ice-cold 1×Native lysis buffer containing 1% NP40 and re-solubilized in 1×Laemmli Sample Buffer (62.5mM Tris-HCl pH6.8, 2% SDS, 10% Glycerol). Protein concentrations were determined by using BioRad DC™ Protein Assay.

### Mass spectrometry

200μg of protein samples from pellet, supernatant or total cell lysate fractions were in-gel trypsin digested as previously described (Shevchenko et al., 1996). The resulting peptides were desalted with high-capacity C18 (Phenomenex) STAGE Tips before an offline high pH reversed-phase chromatography fractionation as previously described (Udeshi et al., 2013). Fractions were collected at 2 minutes intervals. The resulting fractions were pooled in a non-contiguous manner into 8 fractions for SILAC experiment. Each fraction was then dried in a Vacufuge plus (Eppendorf).

Mass spectra were acquired on an Impact II (Bruker) on-line coupled to either an EASY-nLC 1000 (Thermo Scientific) or a nanoElute (Bruker) liquid chromatography (LC). The LC was equipped with a 2-cm-long, 100-μm-inner diameter trap column packed with 5 μm-diameter Aqua C-18 beads (Phenomenex) and a 40-cm-long, 50-μm-inner diameter fused silica analytical column packed with 1.9 μm-diameter Reprosil-Pur C-18-AQ beads (Dr. Maisch). Samples were run with a 120 minutes gradient from 10% Buffer B (0.1% formic acid and 80% acetonitrile in water) to 17% Buffer B during the first 45 min, then Buffer B was increased to 35% by 90 min and to 80% at 95 min. The scanning range was from m/z 200 Th to m/z 2000 Th. The data-dependent auto-MS/MS mode was set to fragment the 17 most abundant ions with the exclusion window of 0.4 minutes.

All raw proteomics data have been deposited to the ProteomeXchange Consortium through the PRIDE partner repository with the identifier PXD012631 (Deutsch et al., 2017; Vizcaíno et al., 2014). The results were analyzed against SGD_orf_trans_all_20110203 released from the *Saccharomyces* Genome Database (yeastgenome.org) with common contaminants with the latest version of MaxQaunt software when the data was generated (version 1.5.2.8). The searches were done using the default software settings, plus match-between-runs and re-quantification and an FDR set below 0.01.

### Bioinformatics

The median log_2_ ratio of total cell lysate was calculated and subtract from all supernatant ratio to correct for error in mixing. Samples were then binned according to loss of solubility upon heat shock or PCC score. Bins were validated by altering the location of bin limits and bin sizes and observed trends in the data did not change (data not shown).

All figures were generated using the R package and assembled using Adobe Illustrator(R Core Team, 2018) Computational analysis tools were carried out using in-house Perl scripts and statistical tests were performed in R. In all cases of non-binary data, p-values were obtained using the Mann-Whitney-Wilcoxon test in conjunction with the Benjamini-Hochberg procedure for multiple testing corrections to determine significance thresholds and reduce the false discovery rate (Bauer, 1972). In the case of binary data, the data was scrutinized against a hypergeometric distribution to determine significance. IDRs were determined with DISOPRED3 (Jones and Cozzetto, 2015). Patches of disordered amino acids where identified using a sliding window of 30 amino acids which allowed for up to 5 amino acids to be considered as ‘not disordered’ before truncating the disordered stretch. Protein abundances were used from the integrated yeast proteome on PaxDb 4.0 (Wang et al., 2015). Solvent exposure calculations were performed using the SSpro8 and ACCpro software in the SCRATCH-1D 1.1 package (Magnan and Baldi, 2014). GRAVY score was calculated with in house scripts using previously derived parameters (Kyte and Doolittle, 1982). Motif identification was carried out using the Analysis of Motif Enrichment (AME) Tool (McLeay and Bailey, 2010) in the MEME-Suite of software (Bailey et al., 2009) using the Fisher test criteria, the average scoring method, and a background of shuffled proteins of the yeast proteome. Low complexity regions were calculated using fLPS (Harrison, 2017) with default settings. Apparent T_m_ data was taken from the study performed by Leuenberger *et al*. (Leuenberger et al., 2017). While RNA interacting proteins where identified from cross-linking studies by Beckmann et al. (Beckmann et al., 2015), Phosphorylation data was obtained from dbPTM (Huang et al., 2016). Molecular recognition features (MoRFs) were identified by the MoRF CHiBi System (Malhis et al., 2015).

Aggregation prone patches and prion-like composition were determined by TANGO (Linding et al., 2004) and PLAAC (Lancaster et al., 2014) software packages, respectively.

### Microscopy

For epifluorescence microscopy, cells were grown to mid-log phase (∼OD_600_ = 0.6) in synthetic defined medium with low fluorescence yeast nitrogen base (loflo, Formedium) and 2% dextrose. 1ml of cultures were heat shocked in a thermomixer at 45°C for 15 minutes. Cells were subsequently collected by centrifugation at 6,000x*g* for 30 seconds. For starvation, cells were incubated loflo medium without dextrose for 90min before imaging. For azide treatment, cells grown at 25°C were treated with 1% NaN_3_ (w/v) for 30 minutes at 30°C before imaging. Live cells were resuspended in growth media before imaging with a Zeiss AxioObserver Z1 equipped with a 470nm and 590nm Colibri light sources and a 63×1.4 NA oil immersion DIC objective. Images were acquired and processed with the Zen2 software.

The diploid strains with Pab1-mCherry and a candidate protein fused to GFP were inoculated into a 384-well glass-bottom optical plate (Greiner) using a pintool (FP1 pins, V&P Scientific) operated by a Tecan robot (Tecan Evo200 with MCA384 head) and grown at room temperature overnight. Upon reaching an OD_600_ equal to ∼0.5 the plate was heated by a plate heater which temperature was set to 48°C (Torrey Pines Scientific) for 20 minutes. Following the heat shock, imaging was performed with an automated Olympus microscope X83 coupled to a spinning disk confocal scanner (Yokogawa W1), using a 60× objective (Olympus, plan apo, 1.42 NA). All strains were imaged within a 15 minutes time period. Excitation was achieved with a green L.E.D for brightfield images, a 488 nm laser (Toptica, 100 mW) for GFP, and 561 nm laser (Obis, 75 mW) for mCherry. Emission filter sets used to acquire the brightfield, GFP and mCherry images were 520/28, 520/28 and 645/75, respectively. The same triple-band dichroic mirror was used for all channels (405/488/561, Yokogawa). Images were recorded on two Hamamatsu Flash4-V2 cameras, one for the brightfield and GFP channels and the second for the mCherry channel. Each image set was composed of two brightfield (BF) images (one in focus and one defocused to facilitate cell segmentation, each with 30ms exposure) as well as one image for each fluorescent channel (500ms exposure for GFP and 500ms exposure for Pab1-mCherry). The focus was maintained throughout the experiment by hardware autofocus (Olympus z-drift compensation system).

To detect colocalization in individual cells, we first segmented cells using the brightfield image, as previously described, and refer to each cell as a region of interest (Matalon et al., 2018). To quantify subcellular colocalization within each cell, we then select pixels with fluorescence intensity above a brightness threshold (30th percentile) and calculated, on those, the pearson correlation coefficient (PCC) between green and red pixel intensities:

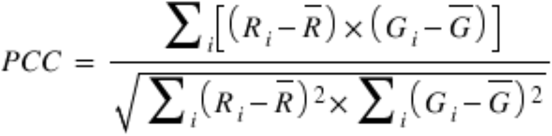

where R_i_ and G_i_ refer to the intensity values of the pixels in the red and green channels, 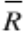 and 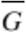 refer to the mean intensities of the red and green channels, respectively, across the entire cell. PCC values range from 1 for cells whose fluorescence intensities are completely linearly related, to −1 for cells whose fluorescence intensities are totally inversely related to one another. The median PCC calculated on a cell population (of typically 300 cells) quantifies the colocalization of a candidate protein with Pab1. All the scripting for image analyses was carried out using Fiji/ImageJ (Schindelin et al., 2012).

### FRAP

Images were acquired using an Olympus Fluoview FV1000 laser scanning confocal microscope with an UplanSApo 100× 1.40 Oil immersion objective. The selected region of interest (ROI) was photobleached with a 405nm diode laser. Images were acquired every 5 seconds for 20 frames after the initial photobleach. For FRAP (Fluorescence recovery after photobleaching) the signal was monitored in the photobleached ROI. For FLIP (Fluorescence loss in photobleaching), the signal of three additional ROIs that contain cytosolic foci was averaged. Three additional ROIs from adjacent cells that were not bleached were also monitored for photofading. Fluorescence of ROI was calculated by first removing background signal, corrected for photofading and normalized across replicates by standardization and feature scaling.

### Western Blotting

100ml culture of cells were either directly collected at OD_600_= 0.8-1 or subjected to heat shock at 45°C for 20min. Cells were washed twice with 1×Native lysis buffer before resuspension in 100µl of 1×Native lysis buffer and lysis with glass beating. Cell debris were then removed by two 1,000×*g* centrifugation steps for 5 minutes. 100µl of total cell lysate was centrifugated at 16,100×*g* for 15 minutes. 25µg of total cell lysate and equal corresponding volumes of supernatant and pellet fractions were resolved by SDS-PAGE using 4%-20% gradient gels for Western blotting with the anti-GFP antibodies (1:1,500 Roche) and LI-COR secondary antibodies (1:10,000). Images were captured and quantified using the CLx Odyssey system using Image Studio v3.1 (LI-COR Biosciences).

## Acknowledgments

We thank Dr. P. Hieter and Tejomayee Singh for sharing strains from the GFP Collection, Dr. N. Stoynov for his help with mass spectrometry experiments, Dr. G. Cohen Freue for advice and members of the Mayor lab for discussions. This work was supported by a grant from the Natural Science and Engineering Research Council of Canada (NSERC). T.M. is the recipient of Career Awards from the Michael Smith Foundation for Health Research and M.Z. is the recipient of the Alexander Graham Bell Canada Graduate Scholarships-Doctoral and the Killam Scholarship.

## Contributions

M.Z. designed and carried out most of the experiments through discussions with T.M.; E.K. carried out the computational analyses with additional participation from M.Z.; J.Z. performed part of the IDR analysis; O.M. carried out Pab1 colocalization and B.D. carried out image processing and PCC calculation; A.H. helped with FRAP data acquisition and analysis; C.L., E.L. and J.G. helped design experiments related to FRAP, co-localization, and computational analyses, respectively. J.Z. and E.K. equally contributed to this manuscript. M.Z. and T.M. wrote the paper and all other authors edited or commented on the manuscript.

---

**Figure S1.**
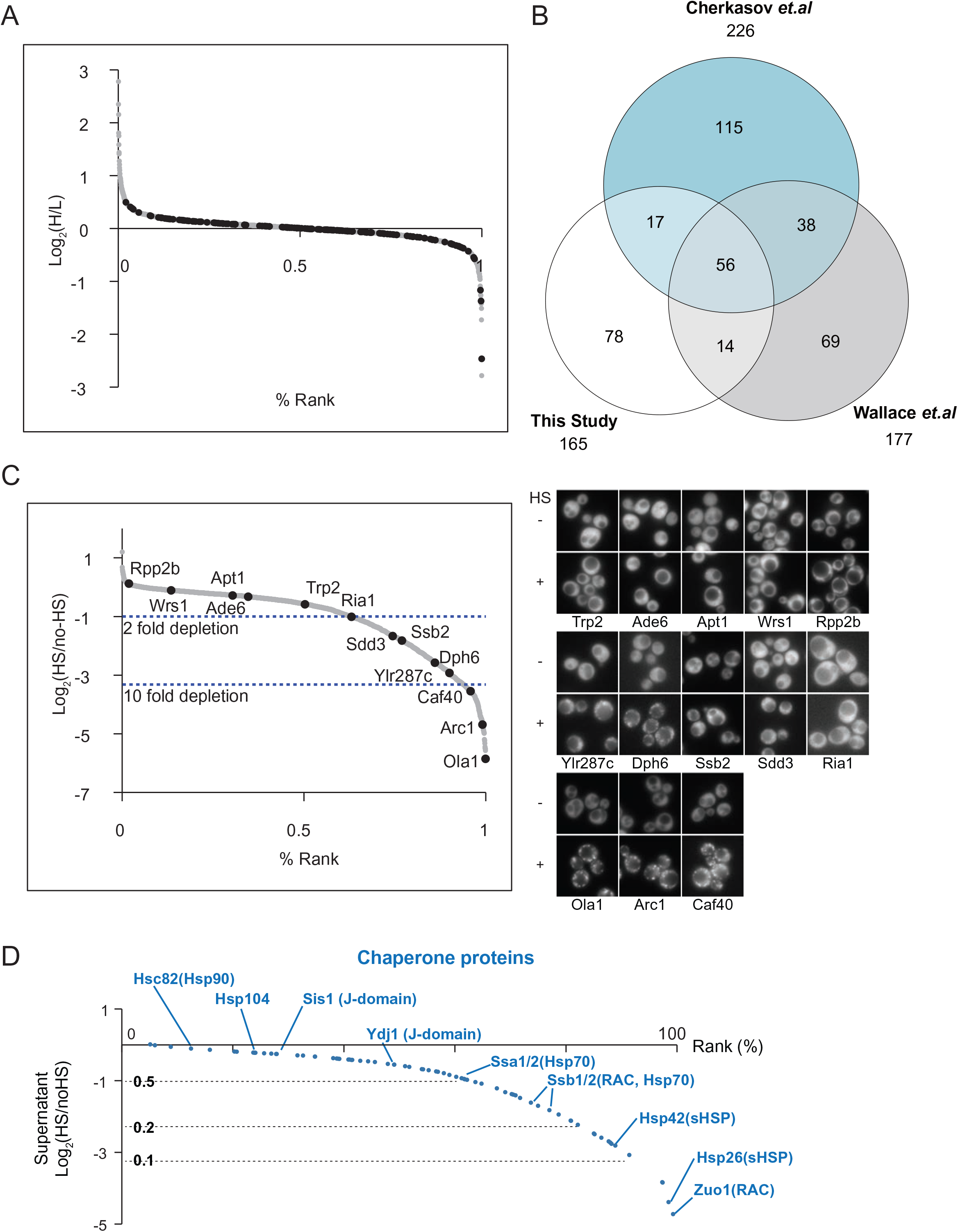
(A) Log_2_ ratios of 3196 proteins after and before heat shock from total cell lysate. Black dots designate proteins that were depleted from both the pellet and supernatant fractions in Figure 1D (data points in the lower right portion of the plot). (B) Comparison of proteins that sediment upon heat shock that were identified in this study and in Wallace *et al*. and Cherkasov *et al*. (Cherkasov et al., 2015; Wallace et al., 2015) (C) Representative images of a panel of cells that were heat shocked for 15 minutes at 45°C (HS) with 13 GFP-tagged proteins randomly selected from the proteomic analysis. The plot in the left shows the averaged log_2_ ratios of the selected proteins in the mass spectrometry analysis. (D) Distribution of 73 quantified chaperone and co-chaperone proteins in the supernatant fraction (blue data points). Names of selected chaperone proteins are shown.

**Figure S2.**
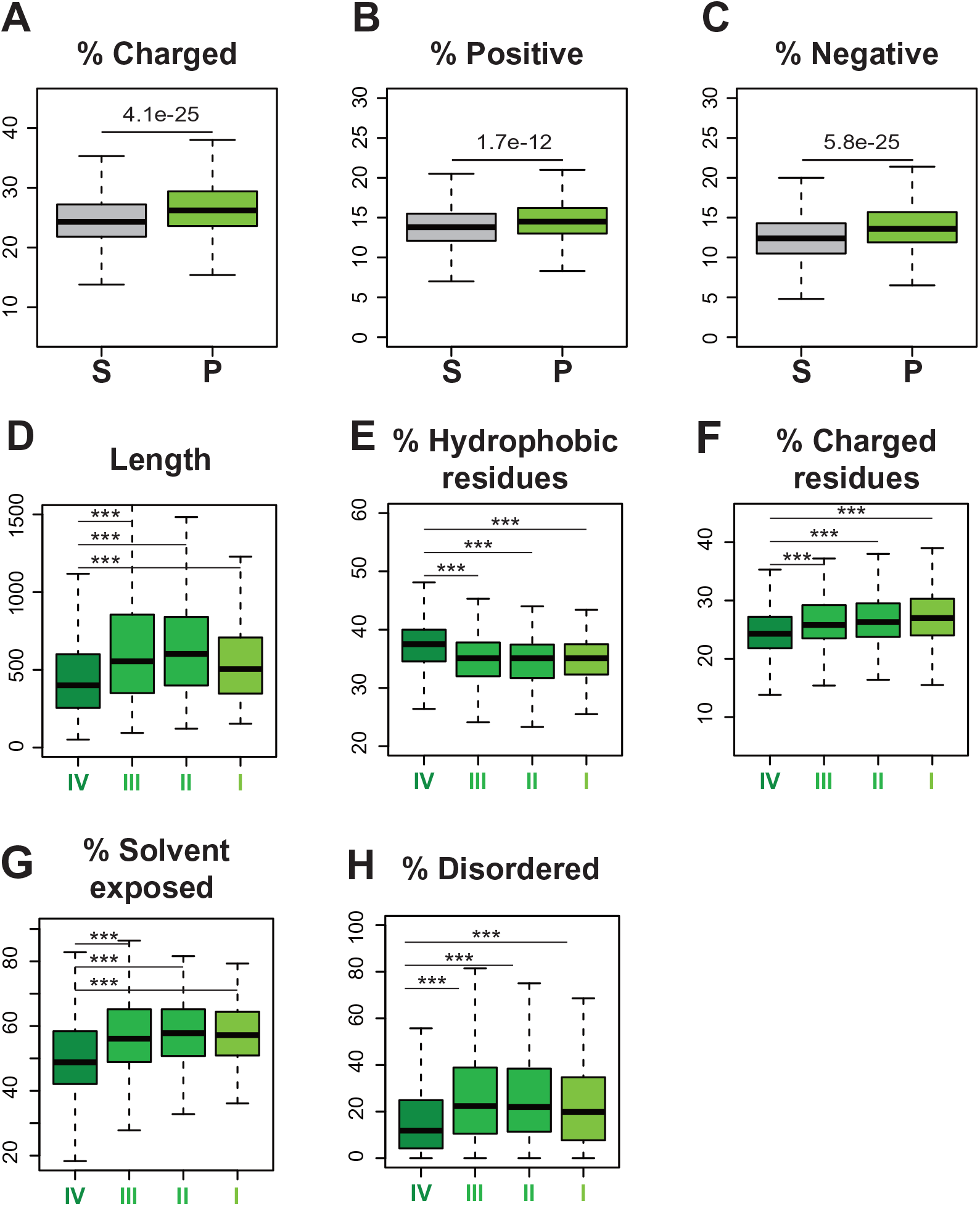
Feature analysis of pelletable proteins upon heat shock. (A-C) Box plots comparing the distributions of the indicated features for soluble (S) and pelletable (P) proteins for % charged residues (A), % positively charged residues (B), % negatively charged residues (C). p-values are indicated. (D-H) Box plots comparing the distributions of the indicated features in the designated bins (see also Figure 2A) for protein length (D), % hydrophobic residues (E), % charged residues (F), % solvent exposed (G) and % disordered (H). All n and p-values are reported in Table S4.

**Figure S3.**
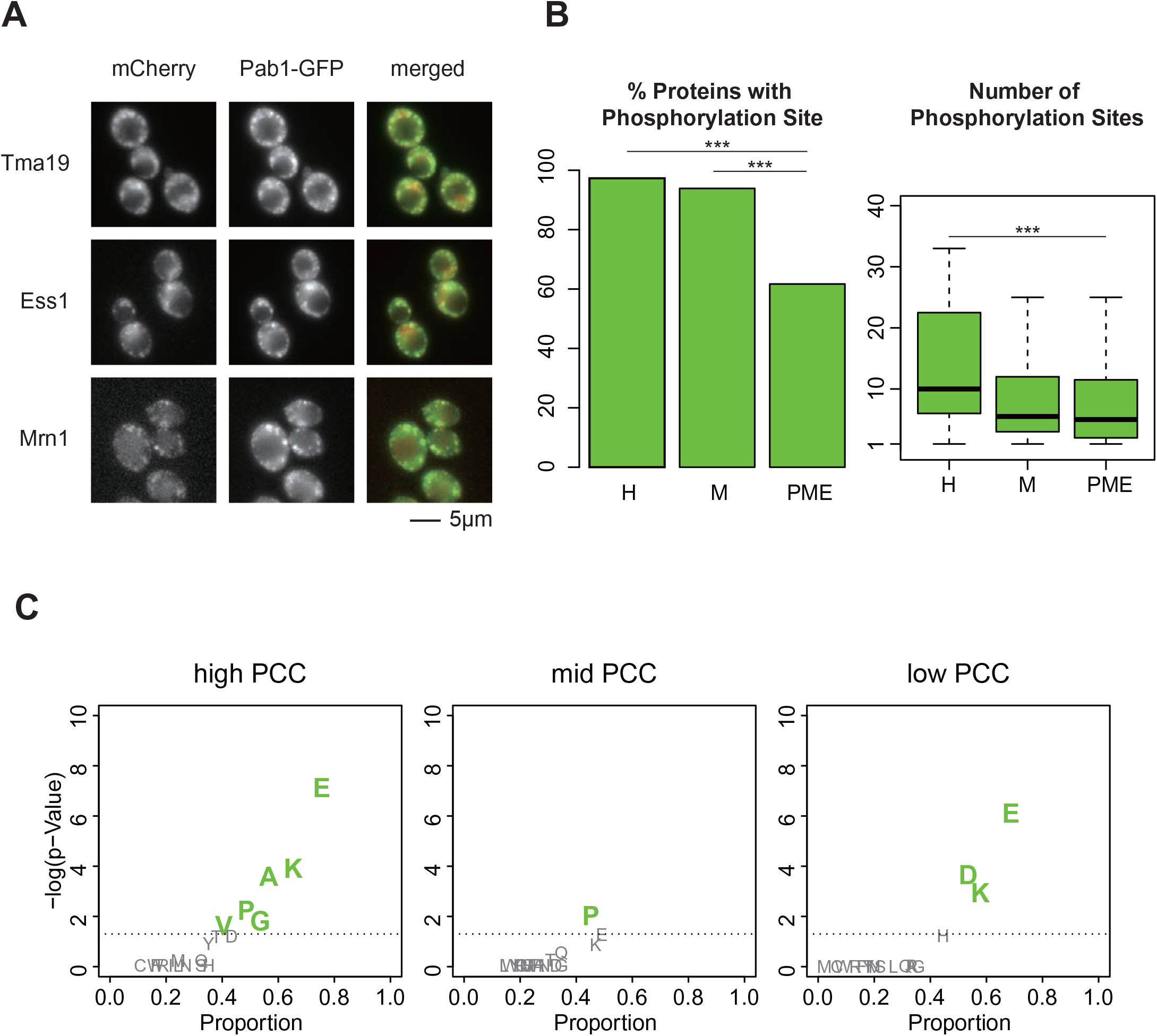
(A) Colocalization of the designated proteins tagged to mCherry Tma1, Ess1, and Mrn1 with Pab1-GFP. (B) Bar plot (left) of the fraction of proteins that was found to be phosphorylated within the proteins with high (H) or mid-range (M) PCC, and in the proteome (PME). Box plot (right) of the distributions of the number of phosphorylation sites among the phosphorylated proteins with high (n = 36) and mid-range (n= 46) PCC or within the proteome (n = 4149). (C) Scatter plots of low probability sequence of a specific amino acid indicating the presence of corresponding LCR. The proportion indicates the portion of proteins containing the corresponding LCR for SG proteins with high (left) and mid-range (center) PCC, and proteins with low PCC values or that did not localize in SG (right). The n and p-values are reported in Table S4. These results indicate that LCR with A, P, G and V were specifically enriched among SG proteins with high PCC values in comparison to highly pelletable proteins not localizing to SG (i.e., low PCC).

**Figure S4.**
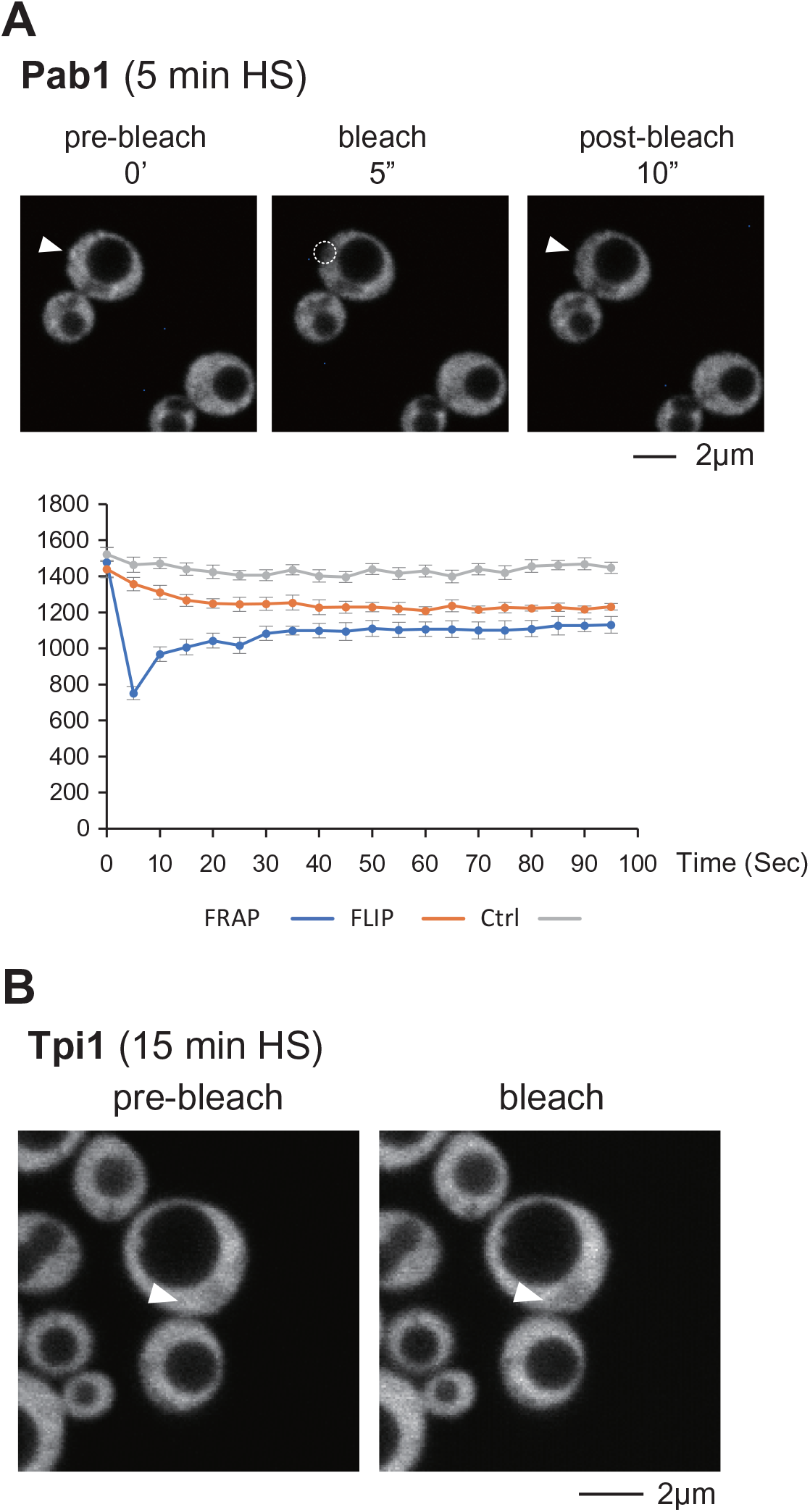
(A) Fluorescent Recovery After Photobleaching (FRAP) of Pab1 after 5min of heat shock at 45°C. The photobleached ROI and the three ROI for FLIP were located in the cytosol as no clear foci could be observed. Intensities from three ROIs in unbleached adjacent cells are also shown (Ctrl). In this particular case, no scaling was applied at time 0, as some of the signal in the photobleached ROI had recovered. The experiment was repeated six times. A representative cell is shown on the right. (B) Image of a Tip1-GFP cell heat shocked for 15min at 45°C before and after photobleaching. The arrows point toward the photobleached ROI that could not be clearly delimited due to the rapid signal recovery.

**Figure S5.**
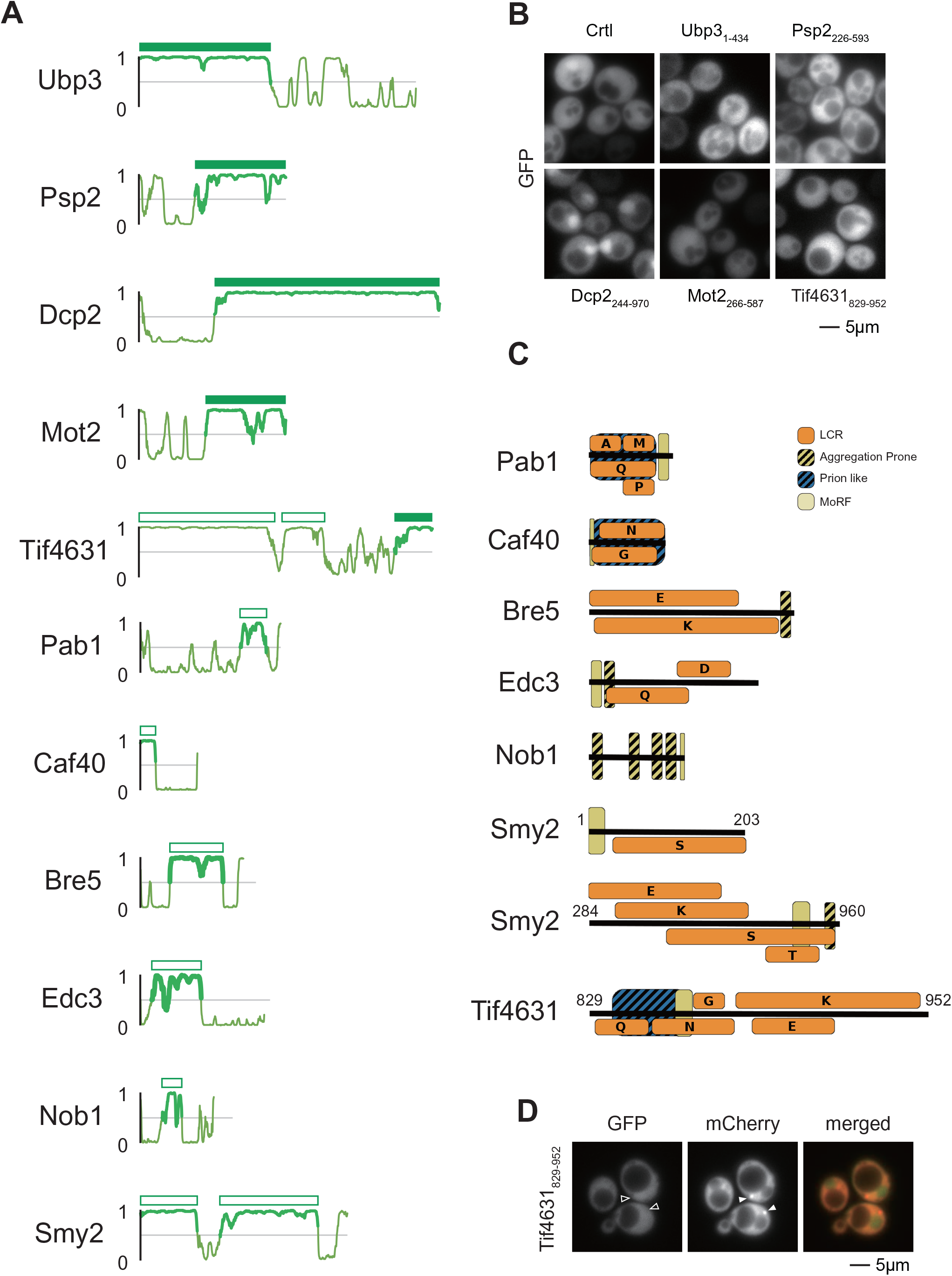
(A) Illustration of predicted disordered score of each tested protein. Bar above the plot shows regions that were fused with GFP and tested for whether they form foci and colocalize with Pab1 under heat shock condition. Solid bars indicate IDRs that colocalized with Pab1 in heat shock induced foci and empty bars designate IDRs that did not localized in Pab1 foci. In the case of Tif4631 and Smy2, multiple IDRs from the same protein were tested. Of note, Bre5 is also a cofactor of Ubp3 that is required for the deubiquitinase activity, and which binds to a Bre5 binding domain within the IDR of Ubp3 (Li et al., 2007; 2005). In contrast to the IDR of Ubp3, Bre5’s IDR could not be directly recruited to SG. (B) Representative unstressed cells (grown at 25°C) expressing the indicated IDR. The images were acquired using the same parameters as in Figure 5A. (C) Feature analysis of the IDRs that were not recruited to stress granules. Orange boxes indicate low complexity regions with letters to indicate specific amino acid enrichment, black striped boxes indicate predicted aggregation prone regions, blue striped boxes indicate prion-like composition, and yellow boxes indicate predicted MoRFs. Only highly significant (p ≤ 1e-6) low complexity regions are pictured. (D) Representative image of Pab1-mCherry cells carrying the designated IDR of Tif4631 after a 90min starvation. Filled arrowheads designate Pab1 foci and hollowed arrowheads their corresponding location in the GFP channel, in which no distinguishable foci is observed.

**Figure S6.**
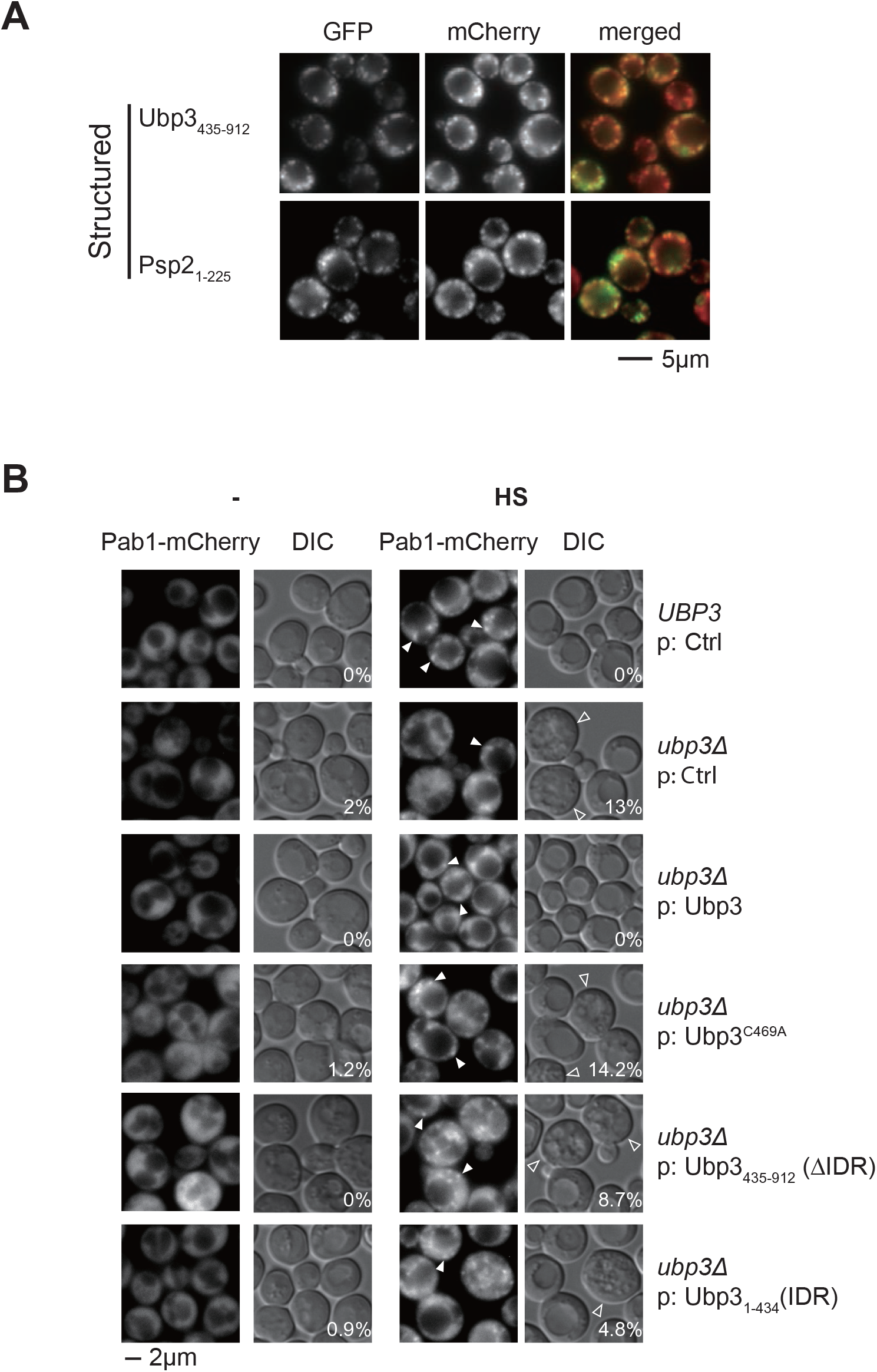
(A) Representative images of colocalization of Pab1-mCherry with truncated Ubp3 and Psp2 that lack their IDR and tagged with GFP after heat shock. (B) Representative DIC and Pab1-mCherry images of cells with the indicated genetic background and protein expressed from a plasmid (p) before and after 15min heat shock (HS) at 45°C. Ctrl designates the control pRS313-EGFP plasmid. Hollowed arrowheads in the DIC images designate cells that lack a distinguishable large vacuole, and the numbers indicate the percentage of such cells (150-300 cells counted per condition; viability of cells lacking a large vacuole was confirmed by methylene blue staining). Arrowheads in Pab1-mCherry images designate representative foci. We could not distinguish a different in numbers of Pab1 foci or the number of cells with Pab1 foci in the tested conditions. Note that, in the previous report, Pbp1 and Pbp4 foci but not Pab1 foci were assessed (Nostramo et al., 2016).

